# State-dependent dynamics of cuttlefish mantle activity

**DOI:** 10.1101/2023.06.19.545578

**Authors:** Sophie Cohen-Bodénès, Peter Neri

## Abstract

Cuttlefish skin is a powerful rendering device, capable of producing extraordinary changes in visual appearance over a broad range of temporal scales. This unique ability is typically associated with camouflage, however cuttlefish often produce skin patterns that do not appear connected with the surrounding environment, such as fast large-scale fluctuations with wave-like characteristics. Little is known about the functional significance of these dynamic patterns. In this study, we develop novel tools for analyzing pattern dynamics, and we demonstrate their utility for detecting changes in feeding state that occur without concomitant changes in sensory stimulation. Under these conditions, we find that the dynamic properties of specific pattern components differ for different feeding states, despite no measurable change in the overall expression of those components. These dynamic changes are therefore not detectable by conventional analyses focusing on pattern expression, requiring analytical tools specifically targeted to pattern dynamics.

## INTRODUCTION

The cryptic abilities of cuttlefish, known as “chameleons of the sea,” are justly celebrated as stunning examples of animal camouflage. Cuttlefish can achieve nearly complete perceptual disappearance against their natural environment (Kelman et al., 2008), and their skin repertoire is able to approximate specific features of artificial backgrounds (Barbosa et al., 2008; Chiao et al., 2010; Osorio et al., 2022). They also engage in complex mimicry, for example by impersonating crabs to the extent of incorporating characteristic locomotory behaviors (Okamoto et al., 2017).

Skin patterning is not restricted to camouflage, however (Hanlon et al., 1988): cuttlefish and other cephalopods produce dynamic patterns in association with specific interactions between conspecifics (Laan et al., 2014; Schnell et al., 2015a;b; 2016; Allen et al., 2017), such as the chromatic pulse pattern displayed by rival male cuttlefish engaged in aggressive or territorial behavior (How et al., 2017), or the passing wave pattern displayed during hunting (Hanlon and Messenger, 2018). It is clear that skin patterning does play a role in some forms of communication that bear no obvious connection with camouflage (Langridge et al., 2007; Galán et al., 2020).

The two functions outlined above, namely camouflage and communication, are generally associated with measurable changes in the *overall expression* of specific skin patterns: for example, when camouflaging against rocks, cuttlefish may display the disruptive pattern more often than other patterns; when camouflaging against sand, they may display the uniform pattern more often. When using the term “overall expression,” we refer to the average intensity of a given pattern over a relatively long period of time, not to the manner in which its intensity changes instantaneously from one time point to the next as time goes by. When referring to the latter characteristic, we use the term “dynamics.” These two characteristics are not necessarily coupled: two patterns may be equally expressed under two different conditions, however they may display different dynamic characteristics under those two conditions. This class of phenomena represents the focus of our study.

## MATERIALS AND METHODS

### Animals and husbandry

We conducted behavioral experiments using 13 adult cuttlefish (Sepia officinalis, 7 females and 6 males) ranging in age between 3 months (∼ 3 cm mantle length) and 12 months (∼ 15 cm mantle length). All animals were reared from eggs and maintained from hatchling following established guidelines (Fiorito et al., 2015), either at our facility in Paris or at the authorized institutional supplier Aquarium de la Rochelle (La Rochelle, France). Eggs were collected from the Atlantic Ocean and cuttlefish were reared in captivity (hatched from eggs) by a team of biologists specialized in cuttlefish breeding. When eggs or juveniles were transported from Aquarium La Rochelle to our facility, transportation was optimized by oxygenation enrichment, noise/vibration minimization in a cool specialized container without light exposure, and following food deprivation before transport to avoid accumulation of toxic ammonia level. Once eggs/juveniles reached our facility, they were acclimatized in three artificial marine water tanks (300 L each) for 2 hours. Their health was closely monitored (every hour) for the subsequent 48 hours, and daily until natural death (we carried out euthanization with a MgCl2 solution whenever necessary). When handling eggs, we introduced additional air pumps to provide sufficient oxygenation for their development.

We ensured that the environment of all three fiberglass tanks approximated the natural habitat as closely as possible, while at the same time providing adequate monitoring of water quality and animal health. Tanks operated a closed water system that allowed constant monitoring and optimization of water parameters, and contained various oxygen pumps, skimmer, mechanical/biological filtration, and carbon filters for ink removal. We introduced sand enrichment (7-cm layer of natural marine uniform sand with granulometry of 1–4 mm, enriched with natural pebbles of different size/color placed randomly across the tank), natural Atlantic ocean shells, and live rocks (∼15 live rocks of different sizes for a total of ∼20 kg per tank) to provide shelter, and more generally to encourage cuttlefish expression of their full behavioral repertoire (burying in sand, establishing territories and dominance hierarchies, protection from lighting within areas shadowed by rocks).

Tanks were visually isolated from one another to prevent cuttle-fish from interacting with animals in other tanks, and tank surface was opacified to minimize mirror reflections. We allowed a maximum of five animals per tank, limited to one individual whenever the potential for aggressive behavior was observed. We established a light/dark cycle of 12-hour/12-hour, and only conducted experiments during the day. Live food was provided three times a day: at 9 am, at 2 pm, and at 8 pm. We never deprived animals, instead food was systematically provided so that cuttlefish could hunt at will (especially during the night) and would not engage in aggressive behavior for food competition. From hatchling to 1 month of age, cuttlefish were fed a mixture of live Artemias and live Mysis. Once they reached 1 month of age, they were fed bigger live marine shrimps (*Palaemonetes vulgaris* provided by Camargues Peches, Aquadip) and live crabs (provided by Aquarium de Paris) especially raised for feeding marine animals reared in captivity.

### Experimental procedure

Our goal was to record cuttlefish behavior under conditions that would approximate their natural spontaneous state. For this reason, we did not interfere with their daily routine and never moved animals to a different tank for testing. Our observational experiments are therefore classed below threshold for explicit ethics approval as expressed in Directive 63/10. We nevertheless seeked and obtained validation by the veterinary official of our Biology Department (IBENS, École Normale Supérieure, Paris, France). When a cuttlefish settled within a specific area of its choice, we fixed an underwater camera (GoPro Hero7) above the tank to record the mantle for ∼2 hours, three times a day, over a period of ∼9 months. Cuttlefish were used to the presence of the camera and did not appear to express signaling patterns in specific response to camera positioning. Experimenters left the room during recording sessions to avoid the emergence of signalling patterns specific to their presence.

### Rationale behind food consumption as experimental protocol

Our protocol was designed to prompt changes of behavioral state with minimal involvement of camouflage and communication. Both behaviors produce sizeable changes in overall pattern expression (Barbosa et al., 2008; Chiao et al., 2010; Osorio et al., 2022; How et al., 2017; Hanlon and Messenger, 2018; Woo et al., 2023), which introduce numerous challenges for unbiased estimation of pattern dynamics *per se* (Satuvuori and Kreuz, 2018). Furthermore, we were specifically interested in whether cuttlefish may alter their behavioral state in the absence of external stimulation as effected by a changing environment or interaction with conspecifics, and whether these alterations may be reflected on their skin by dynamic changes that are not easily detectable as increases/decreases in the overall expression of certain pattern components.

To achieve the above goals, we selected feeding behavior for the following reasons. First, it is relatively easy to define when the animal initiates/completes feeding (see below for how we defined epochs). Second, if we exclude (as we did) the time period during which the prey is introduced into the tank and is visible to the cuttlefish, there is no obvious difference in visual environment between the periods preceding/following food administration and the period associated with food consumption.

With regard to communication with conspecifics occupying the same tank, this behavior is not expected to be selectively engaged (or not) during feeding. Furthermore, the best individual example of state-dependent dynamic signatures comes from a cuttlefish (indicated by square symbols in **Figure 3B**) that never shared her tank with any other animal. Measurements for this individual are more reliable than those obtained from all other individuals, because we were able to carry out more experiments with this animal thanks to its longevity.

### Video segment classification/selection

Each experimental session produced ∼2 hours of continuous video recording, however we typically retained less than half of the recorded frames because the animal was allowed to behave without constraints, and it took a substantial amount of time for the animal to settle into a stable position within view of the camera. Furthermore, various segments were excluded because unsuitable for further tracking/analysis (see below). We extracted segments from the video file and classed them into one of the three main conditions (before, during, and after eating). At the start of this procedure, we provisionally defined the following epochs: the epoch “before” eating started 5 minutes after the beginning of video recording (to exclude initial potential responses to camera placement), and ended right before the introduction of food (live preys) into the tank; the epoch “during” eating started once the prey was fully inside the buccal mass, and ended when the animal stopped producing buccal movements; the epoch “after” eating started 5 minutes after the end of the “during” phase, and continued to the end of the video recording session. After completing this preliminary subdivision of different sections within the movie, we excluded segments from each section according to the following criteria: 1) the animal performed large movements towards other areas of the tank; 2) the animal disappeared from view; 3) the hand of the experimenter was visible, for example during introduction of the prey; 4) live preys were visible around the cuttlefish. In all these instances, segments were excluded to minimize the potential contributions of confounding factors and/or to facilitate subsequent tracking by our algorithm (see further below). Following the above procedures, the allocation of data mass to the three main conditions was 33% before, 25% during, and 42% after eating. The duration of the different epochs for a given experimental session was within comparable ranges across experiments, with lower–upper quartiles of 4–18 minutes (“before”), 4–14 minutes (“during”), and ≈4–11 minutes (“after”).

### Tracking/alignment

We wrote in-house software to identify the position of the animal on every frame, and to align the mantle across frames (see **Supplementary Video**). More specifically, we created labelling software to manually mark the position of the two eyes (indicated by red/green dots in **Figure 1A**) and the tail (rear end of the mantle, aligned with posterior segment of white square in **Figure 1A**) for a small subset (<0.1%) of the frames. We augmented the resulting labelled dataset via rotation/flipping, and trained a modified version of the Mask-RNN algorithm (He et al., 2017) to identify corresponding markers in unlabelled images. The output from this stage was submitted to a regularization algorithm that enforced consistency for labels across frames, such as stable inter-eye distance, correction of label inversion across eyes, and pruning of large sudden displacements. We then extracted from each image a square region with size equal to twice the distance between the two eyes, positioned such that the anterior segment aligned with the two eyes and its middle point matched the middle point between the eyes (see examples in **Figure 1A–F**). We converted the resulting square images to monochrome (average of three color channels), applied left-right symmetry (effectively restricting our analysis to bilateral information), rescaled them to a common size of 32×32 pixels to enforce uniformity across different recording sessions and animal sizes, and aligned them to a common coordinate frame (**Figure 1G–K**). To exclude contributions from elements outside the mantle, we further extracted the central square region of each frame (indicated by blue square outlines in **Figure 1G–K**). This procedure generated a large stack of ∼1.6 million images, each image containing 16×16 pixels. Below we refer to this dataset as the “main” stack.

**Fig. 1.**
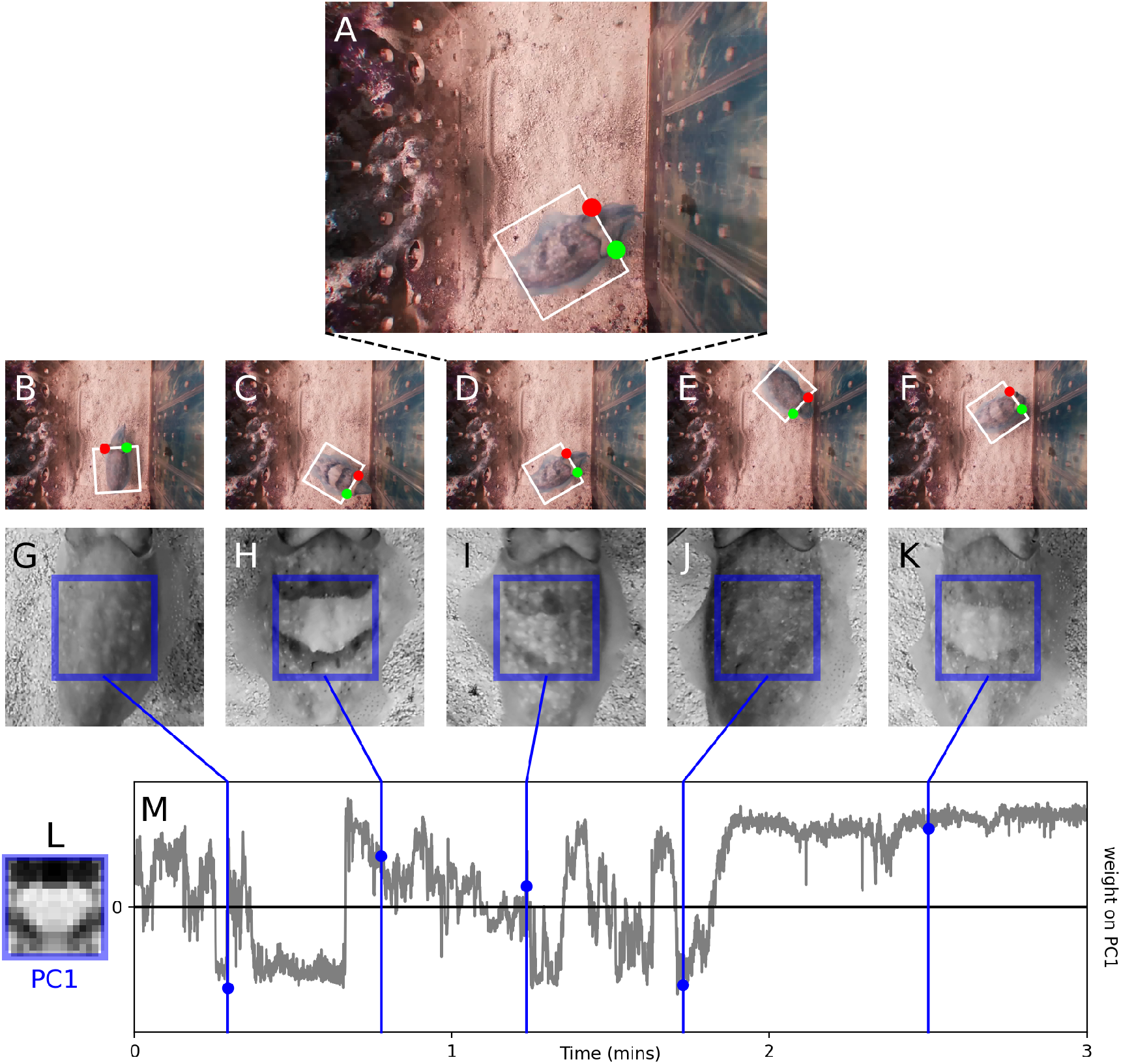
Cuttlefish were filmed from above without disrupting their daily routine/environment (**A**). On each frame (**B**–**F**), an automated algorithm identified the location of the two eyes (indicated by red/green dots) and the full extent of the mantle (indicated by white rectangular outline), which was cropped and rotated to align across frames (**G**–**K**). We applied principal component analysis (PCA) to the sequence of images corresponding to a square region within the mantle (indicated by blue rectangular outlines), and analyzed the time series (**M**) associated with the weight allocated to the first principal component (**L**). Example images in **G**–**K** correspond to time points indicated by blue dots in **M**.

### Principal component analysis

We applied standard principal component analysis (PCA) with varimax rotation to the main stack obtained from tracking/alignment (detailed above). Prior to PCA, the submitted stack was normalized so that each pixel had zero mean and unit standard deviation across frames. In the “aggregate” analysis, PCA was applied to the entire main stack without differentiating across animals (effectively treating the dataset as if it came from one aggregate animal). In the “individual” analysis, PCA was applied separately to sub-stacks obtained by extracting, from the main stack, sections corresponding to individual animals. These two analyses produced nearly identical and internally consistent results (see also further below for quantitative comparisons between individual and aggregate PCs). **Figure 3** shows results obtained using the aggregate approach because it supports ready visualization of each PC in the form of one image that summarizes relevant structure across all animals.

### Overall expression

Each principal component (PC) is associated with a time series recording frame-to-frame weight allocation (see example in **Figure 1M** for the first principal component obtained by applying PCA to a small section of our dataset). We computed the overall weight associated with a given PC by taking the average of the absolute weight values across the time series. When overall weight is computed for each animal and each epoch, it produces an expression matrix of size 13×3 for each PC. The analysis in **Figure 2** shows results obtained using the simple metric described above, however similar results are obtained when using other related metrics, such as standard deviation (SD) across the series.

**Fig. 2.**
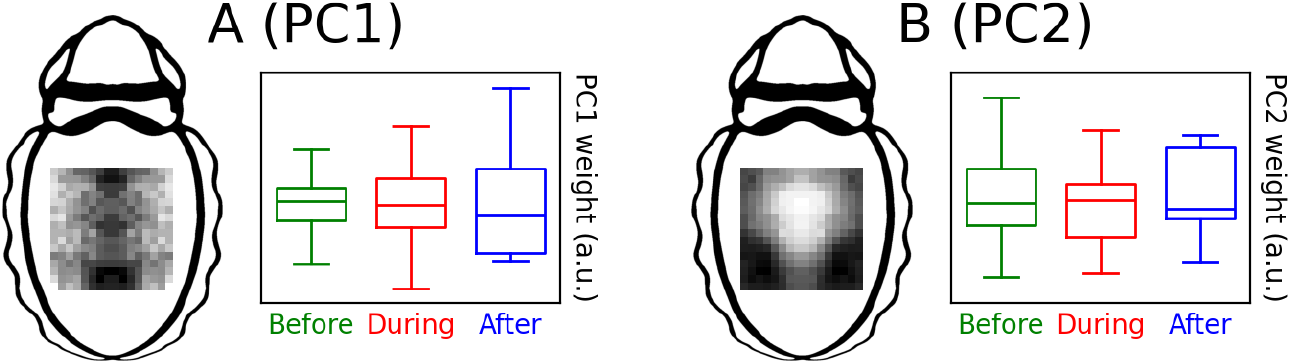
Overall expression of principal components (PCs) does not differ across feeding conditions. The y axis in **A** plots the mean absolute weight allocated to PC1 (shown by image on the left) for the three feeding conditions (before/during/after food consumption indicated by green/red/blue respectively) using conventional box/whisker plots: the box extends from the first quartile to the third quartile of the population (across animals) with a line at the median, while the whiskers extend from the box to the farthest data point lying within 1.5× the inter-quartile range. **B** plots equivalent data for PC2. Units are arbitrary, because absolute units do not carry meaningful information (they depend on how images were rescaled and encoded). The relevant factor in these plots is the comparison across conditions (non-signficant at p=0.79 for **A** and p=0.40 for **B**), regardless of absolute units.

**Fig. 3.**
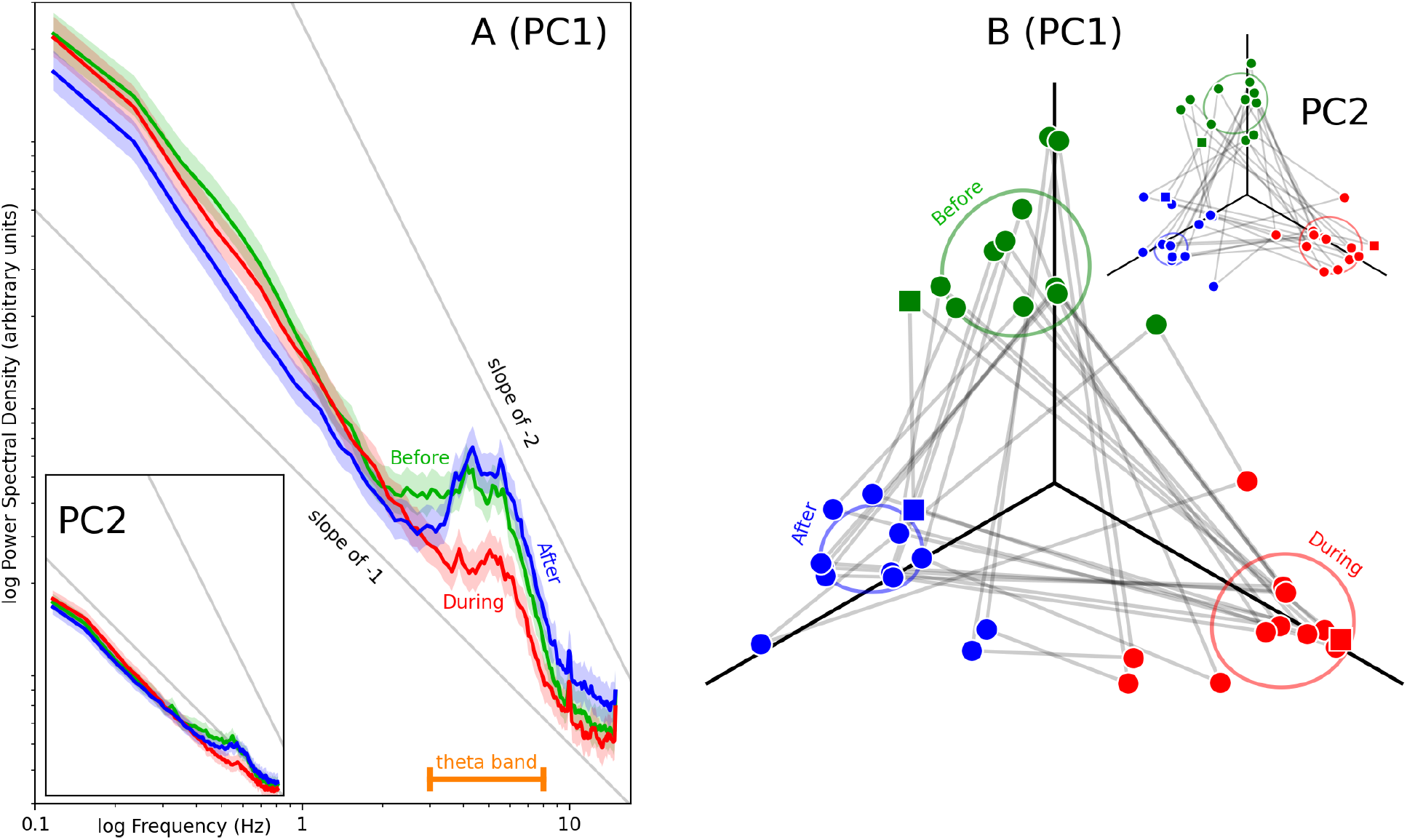
Spectral density profiles for time series (such as the example in **Figure 1M**) associated with weight allocation to PC1, analyzed separately for time periods preceding (green), during (red), and following (blue) food consumption (**A**). Shaded regions indicate ±1 SEM. The dissimilarity between two spectral profiles can be quantified via spectral distance (see Methods), indicated by the length of gray segments joining two dots in **B**. Spectral distance is a unitless normalized quantity that quantifies the difference between two spectral profiles such as those shown in **A**. It is therefore only interpretable when comparing its values across conditions (significant at p<0.01 for PC1, not significant at p=0.58 for PC2), and not with regard to its absolute values for a given condition. This is achieved by comparing the relative lengths of three segments sharing the same gray level, or by comparing relative distances between clusters. Each dot refers to a different animal (n=13) engaged in one of the three feeding conditions (same color convention as in **A**). Square symbols refer to an individual who did not share the tank with any other animals. To aid visualization, dots have been positioned to retain cluster proximity with the three axes defined by black solid lines (see Methods). Oval contours reflect data spread. Insets plot equivalent results for PC2.

### Spectral profiles and distance metric

We computed the power spectral density (PSD) of the time series using Welch’s algorithm as implemented by the scipy package (scipy.signal.welch) with default parameters (in particular, segment length is set to 256 by default), separately for each main condition (before, during, and after eating). As customary when evaluating spectral density, we logged the resulting values (see examples in **Figure 3A** for the first principal component obtained from applying PCA to the entire main stack).

To compute the spectral distance between two vectors of logged PSD values such as, for example, the vectors detailing values for two of the three traces in **Figure 3A**, we first normalized each vector to root-mean-square (RMS) equal 1, and then computed the RMS distance between the normalized vectors. We computed all three possible paired distances for the three main conditions, separately for each animal: distance between before and during, between before and after, and between during and after. To render this metric comparable across animals, we normalized the three values by dividing them by their mean for each animal. When computed across all 13 animals, this procedure generates a distance matrix of size 13×3.

To visualize the distance matrix in **Figure 3B**, we first defined a triangle for each animal with vertices labelled by main condition (before/during/after) and side lengths equal to the three distance values for that animal. We then computed its center of mass, and fixed it to the origin of the three-axis plot in **Figure 3B**. We rotated the triangle around this center to minimize the angular distance between its medians and the three axes of the plot (solid black lines oriented at 60 degrees from one another), with distance calculated between the median to the “before” vertex and the vertical axis, between the median to the “during” vertex and the axis down to the right, and between the median to the “after” vertex and the axis down to the left. This procedure implements clustering of the three vertex types along the three axes, while preserving normalized distances between vertices.

### Correlation between individual and aggregate PCs

To evaluate the correspondence between PCs from individual animals and those from the aggregate animal, we computed the pixel-by-pixel correlation between PCs. When computing a correlation of this kind, it is necessary to accommodate the fact that the overall sign of a given PC is arbitrary. To achieve this goal, we first computed the RMS of the difference between the two PCs to be correlated as returned by PCA, and the RMS of this difference when inverting the sign of the individual PC. If the former RMS value was smaller than the latter, we did not apply sign inversion; if otherwise, we inverted the sign of the individual PC before computing the correlation between the two PCs.

When we quantified the pixel-by-pixel correlation between PCs extracted from the entire dataset and equivalent components extracted from individual animals using the above procedure, we found that the aggregate PC1 obtained from the entire dataset (**Figure 2A**, left) correlates with PC1 estimates from individual animals (correlation values greater than 0 at p×0.001 on a Wilcoxon test), but does not correlate with individual PC2 estimates (p=0.94). The aggregate PC2 (**Figure 2B**, left) correlates substantially with individual PC2 estimates (average coefficient of 0.48, correlation values greater than 0 at p< 0.001). This component also correlates, to a lesser extent (average coefficient of 0.28), with individual PC1 estimates (correlation values greater than 0 at p< 0.01), indicating weaker correspondence between aggregate and invidual estimates for PC2. Overall, this pattern of results is indicative of consistent structure between PC estimates from the aggregate dataset and corresponding estimates from individual animals, with some residual misalignment for PC2.

### Statistical analysis

Our main analysis involved comparison of overall expression and spectral distance across the three main conditions (before/during/after) to confirm the significance (or lack thereof) of the data pattern suggested by the visualizations in **Figure 2** and **Figure 3B**. To achieve this goal, we first submitted the expression/distance matrix to a Friedman test for repeated samples, which returned p=0.79 for PC1 expression (**Figure 2A**), p=0.40 for PC2 expression (**Figure 2B**), p<0.01 for PC1 spectral distance (**Figure 3B**), and p=0.58 for PC2 spectral distance (inset to **Figure 3B**). To determine which conditions differed and which ones did not with regard to spectral distance, we then carried out a posthoc Conover Friedman Test with correction for multiple comparisons (two-stage false discovery rate), which returned no difference (p=0.23) between the before-during distance (separation between green and red clusters in **Figure 3B**) and the during-after distance (separation between red and blue clusters), a clear difference (p<0.01) between the before-during distance and the before-after distance (separation between green and blue clusters), and a measurable (albeit less robust) difference between the during-after distance and the before-after distance (p<0.05).

## RESULTS

Our results can be summarized as follows. We extract principal components (PCs) associated with skin pattern changes (**Figure 1**) during three epochs involving different behavioral states: the epoch preceding the administration of food, the epoch during which the animal engages in food consumption, and finally the epoch following food consumption. We find no changes in the overall expression of specific pattern components across epochs (**Figure 2**), however we do find measurable changes in pattern dynamics (**Figure 3**). Below we describe these results in greater detail, and interpret their significance in the Discussion section.

### Principal component decomposition of skin pattern expression/dynamics

We focus on the dynamics of large-scale patterns across the body of the cuttlefish (Reiter et al., 2018). Because these patterns must be studied with reference to the coordinate frame defined by the body of the cuttlefish, we tracked the animal over time (**Figure 1B–F**) and aligned skin patterns from different time points to form a sequence of co-registered square images (**Figure 1G–K**), which were then subjected to conventional principal component analysis (PCA) to extract the most prominent 2D skin motifs (see Methods for details; see also Kelman et al. (2008); Orenstein et al. (2016); Woo et al. (2023) for related approaches).

**Figure 1L** shows the first principal component extracted from a brief period of a few minutes, alongside the change in weight associated with this component over time (**Figure 1M**). It is evident that the dynamic profile for this component is rich in structure, displaying both quick fluctuations (first 2 minutes of example in **Figure 1M**) and sustained behavior (last minute). In the following, we seek to determine whether this structure undergoes state-specific changes when animals alter their feeding state.

The contribution of a given component can be assessed using different metrics. In this study, we focus on two metrics that capture two fundamentally distinct aspects of the weight contribution associated with different components: *overall expression* (time average of absolute weight values in **Figure 1M**), which measures the extent to which a given component is expressed across a given epoch without regard for its dynamic structure; and *spectral distance* (see Methods), which attempts to estimate dynamic changes associated with component expression between epochs. To clarify this distinction, we may consider a flickering uniform pattern that only takes two intensity values: 0 (dark) and 1 (bright). Over time, the pattern is bright on 50% of the frames, however this may happen in two different ways: in configuration A, the pattern switches color every two frames: 0-0-1-1-0-0-1-1 and so on; in configuration B, the pattern switches color every frame: 0-1-0-1-0-1-0-1 and so on. In both cases, the average overall expression of the bright pattern is 50%, so overall expression cannot discriminate between A and B. However, the difference in dynamic structure (every two frames in A versus every frame in B) can be exposed by measuring spectral distance.

### Overall expression does not change with feeding state

**Figure 2A** plots overall expression of PC1 across the three conditions associated with feeding: before (green), during (red), and after (blue). At this level of analysis, there is no statistically significant difference across conditions (p=0.79). Similar results are obtained for PC2 (p=0.4). We made several attempts at measuring pattern expression using different metrics to identify reliable markers of the feeding state engaged by the animal, but all our attempts failed statistical significance. We conclude that overall expression does not change reliably with feeding state, as may be expected (for example) when comparing different states of camouflage.

### Feeding is associated with reduced dynamics within the theta frequency range

The most obvious tool for summarizing the dynamic range exhibited by **Figure 1M** is spectral density across temporal frequency, as shown in **Figure 3A**. This figure shows spectral density for a single component (plotted in the inset against the body of the animal) extracted from our entire dataset. Weight profiles for this component were analyzed separately for the three conditions associated with feeding (different colors, same color-coding as in **Figure 2**). This analysis overlooks potential differences across animals, in that we are treating data from different cuttlefish as if it were collected from one “aggregate” animal, however our results remain consistent when taking into account individual differences (see **Figure 3B**, Methods, and analysis below).

When comparing spectral density profiles across feeding configurations, we notice a few obvious features. First, the two profiles associated with time periods outside feeding (before and after, green and blue) are very similar. More specifically, the spectral slope of the aperiodic component takes values between -1 (pink noise) and -2 (Brown noise), which is the range typically spanned by many biological signals like the electroencephalogram (EEG) (Pathania et al., 2021), and the periodic component is selectively enhanced within the theta range (Karakaş, 2020) (3–8 Hz indicated by the orange horizontal marker in **Figure 3A**), which presents increased power for both conditions. Second, the profile associated with the feeding period (red) matches the other two profiles with relation to the aperiodic component, however it presents a marked reduction of the periodic component: the power bump within the theta range is almost entirely suppressed.

Based on the qualitative observations outlined above, we devised a metric to quantify the similarity/dissimilarity between two spectral profiles (see Methods). This quantity, which we term spectral distance, can be computed for individual animals and therefore used to create a sufficiently rich dataset to support statistically informed conclusions. **Figure 3B** plots spectral distance in the form of distance across the plane: the length of a segment connecting two data points reflects the spectral distance between the two profiles associated with those points. Each point refers to one animal engaged in one feeding configuration. Overall, this plot confirms our prior observation: data clusters for the two nonfeeding configurations (green/blue) are closer to each other than they are to the cluster for the feeding configuration (red). This pattern is statistically significant for PC1 (see Methods), but not for PC2 (inset to **Figure 3B**).

## DISCUSSION

Our main result is that specific behavioral states experienced by cuttlefish may not be reflected by increases/decreases in the expression of mantle patterns, however they may be associated with specific changes in the *dynamics* of those patterns. More specifically, we found that theta fluctuations of the first principal component extracted from a square region of the mantle is engaged primarily during non-feeding behavior, being almost entirely suppressed during feeding behavior. Below we discuss relevant connections with existing literature, unaddressed interpretational difficulties, and potential implications for our broader understanding of spontaneous pattern dynamics in cephalopods.

### Relations to existing literature

From a behavioral standpoint, relevant literature on cuttlefish behavior during food consumption (the focus of this investigation) is relatively limited. Previous studies on cuttlefish have explored skin pattern changes associated with hunting (Holmes, 1940; Messenger, 1968; Adamo et al., 2006; Cole and Adamo, 2005) and prey catching (Wilson, 1946; Duval et al., 1984; Kim et al., 2022). One prior study indirectly touched on cuttlefish behavior during food consumption (Crook et al., 2002), however it did not examine potential differences in body pattern expression/dynamics during this behavior.

From a technical standpoint, our study shares elements with a previous investigation (Kim et al., 2022) that studied spatial granularity of aligned square regions from the cuttlefish mantle during prey capture. This prior study examined potential changes of spectral energy within separate spatial frequency bands over time, and used those changes to identify different stages during prey capture behavior. While our method similarly relies on extraction/alignment of square regions within the mantle, it differs from this prior study in several important respects (discussed immediately below).

Our approach exposes a novel characteristic of mantle activity that is informative about behavioral state: spontaneous pattern dynamics as a source of information that is separate from, and orthogonal to, pattern expression. The class of pattern changes detected by spatial granularity analysis (Kim et al., 2022) would translate into increased/decreased expression of specific PCs during prey catching, and would therefore be classed as changes in overall expression within the context of our study (**Figure 2**). Our data-driven PC-based spectral analysis (**Figure 3**) specifically targets dynamic features above and beyond pattern expression. In this study, we not only show that these dynamic features carry structure, but also that they carry behaviorally relevant structure.

Furthermore, our methods (from tracking to image metric assessment) are entirely automated. This characteristic removes potential bias associated with manual alignment in previous work (Kim et al., 2022) and provides enhanced scalability, being applicable to larger datasets (such as ours).

### Tentative interpretation with reference to reward-based behavior

Previous research has demonstrated that unexpected food rewards can modify subsequent cuttlefish behavior (Chung et al., 2022), suggesting that changes in internal state may shape foraging strategy for additional food items. We favour an interpretation along similar lines for our own results, namely we hypothesize that feeding-specific changes in pattern dynamics reflect a positive reaction to food consumption, possibly associated with the release of specific neurotransmitters (Heymann et al., 2020; Palovcik et al., 1982). Under this interpretation, the effects we report here may not be specific to food, but may generalize to other forms of reward, including those potentially associated with mating and socialization. In those contexts, it may be more difficult to experimentally isolate skin pattern components that are specific to reward-driven states as opposed to outward-directed communication signals, however similar mechanisms may be at play.

Following on from the above hypothesis, it is tempting to draw a parallel between our finding of theta-specific dynamic changes in pattern expression (**Figure 3A**) and the established role of theta oscillations in reward-based learning across vertebrates (Zold and Hussain Shuler, 2015; van Wingerden et al., 2010; Knudsen and Wallis, 2020), including humans (Lin et al., 2022). A parallel of this kind, however, relies on numerous (and questionable) assumptions, for example that theta oscillations in neuronal activity (such as those that may be measured via local field potentials or EEG (Karakaş, 2020)) are transparently connected with skin dynamics. It is known that specific pattern dynamics are likely controlled by central pattern generators (Laan et al., 2014), and those same generators may also drive EEG signals (Berkowitz, 2019), however there is no evidence to suggest that one may directly compare frequency bands between brain activity and skin dynamics. Furthermore, most evidence on the role of theta oscillations in reward-based and memory-based behavior comes from vertebrates (Karakaş, 2020). For the above reasons, specific interpretations of theta activity in our results remain highly speculative at this stage.

### Internal states in invertebrates?

We conclude by considering the more general question of whether it is plausible to hypothesize that cuttlefish may experience what are termed “internal states” in the literature (Flavell et al., 2022), and whether these animals may express such states via overt mantle displays. It is currently undisputed that invertebrate behavior is conditional upon internal states (Brembs, 2013; Anderson, 2016; Vogt et al., 2021): for example, the satiation state of the predatory sea-slug *Pleurobranchea* regulates approach/avoidance behavior in this animal (Hirayama and Gillette, 2012), and feeding state impacts navigational choices in flies (Vogt et al., 2021), bees (Dyer et al., 2002), and ants (Harris et al., 2005).

It remains unclear, however, whether internal states may also be expressed by skin pattern changes (in animals that are capable of producing them) that are not outward-directed in the immediate sense (Flavell et al., 2022), that is to say not directed towards the environment (camouflage) or other animals (communication). In some lizard and amphibian species, for example, exposure to stressful conditions can dynamically modify body coloration (Boyer and Swierk, 2017; Greenberg, 2002; Kindermann et al., 2013). In all these cases, as in ours, it is difficult to be certain that the observed changes do not reflect some form of communication/camouflage, if not actualized at least intended. Various aspects of our protocol design suggest that camouflage and communication were unlikely to play a role in our experiments (see Methods). While we cannot completely exclude a role for these behaviors, it should be noted that this issue does not impact our primary finding that changes in pattern dynamics may reflect changes in behavioral states that are not measurable at the level of pattern expression.

Notwithstanding the above interpretational difficulties, which are not specific to our study but rather impact all research on internal states (Flavell et al., 2022), based on our results we propose that the possibility of implicit (non-communicative) displays of internal states in cephalopods should be entertained by future studies. The implications brought about by this possibility are wide-ranging (Anderson, 2016; Flavell et al., 2022): from a behavioral standpoint, the existence of such phenomena would provide experimentalists with novel ways to access the cognition of creatures that are evolutionarily distant from, yet possibly share some primordial internal states with, our own species.

